# Primate simplexviruses differ in tropism for macaque cells

**DOI:** 10.1101/2022.07.18.500402

**Authors:** Heike Hofmann-Winkler, Abdul Rahman Siregar, Ignacio Rodriguez Polo, Nesil Esiyok, Sabine Gärtner, Rüdiger Behr, Stefan Pöhlmann, Michael Winkler

## Abstract

Primate simplexviruses are closely related neurotropic herpesviruses, which are largely apathogenic in their respective host species. However, cross-species transmission of Macacine alphaherpesvirus 1 (McHV1, also termed Herpes B virus) from rhesus macaques to humans can cause fatal encephalomyelitis. In contrast, closely related viruses, such as Cercopithecine alphaherpesvirus 2 (CeHV2, also termed simian agent 8) or Papiine alphaherpesvirus 2 (PaHV2, also termed herpesvirus papio 2), have not been linked to human disease and are believed to be largely apathogenic in humans. Here, we investigated whether McHV1, PaHV2 and CeHV2 differ in their capacity to infect non-human primate (NHP) and human cells. For comparison, we included the human simplexviruses HSV1 and HSV2 in our analyses. All five viruses replicated efficiently in cell lines of human and African green monkey origin and McHV1 and PaHV2 also showed robust replication in rhesus macaque cell lines. In contrast, replication of HSV1, HSV2 and CeHV2 in cell lines of rhesus macaque origin was inefficient. These results demonstrate a previously unappreciated partial resistance of certain rhesus macaque cell lines to HSV1/HSV2/CeHV2 infection and reveal similarities between cell tropism of McHV1 and PaHV2 that might be relevant for risk assessment.

## Introduction

Simplexviruses of primates co-evolved with their respective hosts [1, 2]. They share a common genome structure, which is essentially collinear with human Herpes simplex virus type 1 (HSV1; Human alphaherpesvirus 1), the best characterized species of this group. Several simplexviruses from non-human primates (NHP) have been isolated, including Macacine alphaherpesvirus 1 (McHV1, herpes B virus) [3], Cercopithecine alphaherpesvirus 2 (CeHV2, simian agent 8) [4], Papiine alphaherpesvirus 2 (PaHV2, herpesvirus papio 2) [5] and Panine alphaherpesvirus 3 (chimpanzee herpesvirus) [6]. Genome sequencing revealed conservation of all genes among those viruses, with the notable lack of the RL1 (γ34.5) gene in the genomes of McHV1, CeHV2 and PaHV2 [5, 7-10].

The biology of simplexvirus infection of NHP is believed to be similar to infection of humans with HSV1, with a largely asymptomatic primary infection followed by lifelong viral latency in sensory neurons and occasional lesions due to reactivation [11]. This notion is mostly supported by studies analyzing McHV1 infection of macaques kept in captivity [12, 13]. In addition to intraspecies transmission, cross-species transmissions have been frequently documented, especially when different NHP species were cohoused. In many cases, such transmission events (e.g. for McHV1) have been recognized because of apparent or fatal disease [14, 15], but asymptomatic infections have also been documented [16]. Notably, transmission of McHV1 from rhesus macaques to humans leads to encephalomyelitis with high case-fatality rate [17]. In contrast, transmission of CeHV2 and PaHV2 to humans has not been reported, despite these viruses being 79-86% identical to McHV1 on the genome level [10] and it is the general assumption that these viruses do not cause disease in humans.

Cell culture and animal studies highlight the potential of primate simplexviruses for cross-species transmission. Thus, HSV1 has been reported to replicate in cell lines from species as diverse as human, NHP, hamster and mouse [18]. In addition, several mammalian species served as animal models for primate simplexviruses. Infection of mice is a common model to study neuropathogenicity of primate simplexviruses [19-22]. In addition, rabbits and guinea pigs have been employed to study latency by HSV1 and McHV1 [23, 24]. Only recently the use of rhesus macaques as animal model for HSV1 [25-28] and HSV2 [29] has been reported. In contrast, older studies did not detect appreciable replication of HSV2 in macaques [30] and reported poor replication of HSV1 and HSV2 in macaque cell lines [31-33].

We investigated whether the presumed differential pathogenicity of CeHV2/PaHV2 and McHV1 for humans correlated with their capacity to infect cell lines of NHP and human origin.

## Materials and Methods

### Cell culture

Cell lines 293T (DSMZ ACC 635) [34], A549 (ATCC CCL-185) [35], U251 (U373-MG) (ATCC HTB-17; kind gift by T. Stamminger) [36], LLC-MK2 (ATCC CCL-7) [37], sMAGI (NIH ARP5033) [38], TeloRF (kind gift by S. Voigt) [39], Vero76 (ATCC CRL-1587; kind gift by A. Maisner) [40] and Cos7 (ATCC CRL-1651) [41] cells were cultivated in DMEM supplemented with 10% FCS and Pen/Strep. Human cell lines were authenticated by STR typing following a published protocol [42]. The species identity of primate cell lines was authenticated by sequencing part of the mitochondrial CytB gene after PCR amplification [43].

Rhesus macaque induced pluripotent stem cell lines (iPSC lines) used as input cells for the neural aggregates were previously generated and maintained according to Stauske at al [44]. Characterization of the pluripotent state and identity of iPSC lines was performed routinely.

### Rhesus macaque neural aggregate generation and culture

Neural aggregates were generated in stationary conditions following a protocol adapted from Lancaster et al. and Mansour et al. [45, 46]. For generating neural aggregates, iPSCs were dissociated to single cells using Accutase, and 10,000 cells per well were transferred in 96-well ultra-low attachment plate in UPPS culture medium [44]. The medium was supplemented with 5 μM of pro-survival compound (ROCK2 inhibitor; Calbiochem DDD00033325) for the first 24 hours. On day 3, embryoid bodies were transferred to Neural Induction Medium (NIM) (DMEM/F-12 (1:1), N2 Supplement, 20% KnockOut Serum, 3% Fetal Bovine Serum, 1% non-essential amino acids, 2 mM GlutaMAX). The NIM was supplemented with 1 μg/ml heparin, 200 μM L-Ascorbic acid, 10 ng/ml of bFGF2, 10 μM SB431542, 2.5 μM dorsomorphine, and 1 mM sodium pyruvate for the first 4 days; and without heparin, bFGF, and sodium pyruvate for the subsequent 4 days with medium change every other day. On day 11, neurospheres were embedded in Matrigel (10 mg/mL). After removing the culture medium, 50 μL of Matrigel drops were added on top of neurospheres and allowed to polymerize for 20 minutes at 37°C. After incubation, the Matrigel-embedded neurospheres were transferred to 48-well plates coated with anti-adherent rinsing solution in cerebral differentiation medium I (CDM I) (DMEM/F12: Neurobasal Medium (1:1), N2 supplement, B27 supplement without vitamin A, 1% non-essential amino acids, 2 mM GlutaMAX, and 2.8 ng/mL insulin). The CDM I was supplemented with 20 ng/mL bFGF and 20 ng/mL EGF, and the medium was changed every other day. After 5 days, B27 supplement was added for the subsequent 7 days (CDM II) with medium change every other day. From day 22 onwards, EGF and FGF2 were replaced with 20 ng/mL BDNF and 20 ng/mL NT3 (CDM III). The medium was changed every other day. After generation, neural aggregates were used for characterization and infection experiments between days 70 and 100 of differentiation.

### Viruses

HSV1 strain 17syn* and HSV2 strain 333 were a kind gift by Wali Hafezi (Institute of Virology, University Hospital Münster). CeHV2 (SA8) strain B264 and PaHV2 (HVP2) strain X313 were a kind gift by David Brown and Matthew Jones (Public Health England). McHV1 (herpes B virus) was a kind gift of Christiane Stahl-Hennig (German Primate Center). Viruses were propagated on Vero76 cells by infection at MOI 0.01 and harvested after complete cytopathic effect had developed.

### Viral replication kinetics and titration

For one-step growth curves, Vero76 cells were seeded in 24 well plates at 60,000 cells/ml. On the next day, cells were infected with MOI 1 of respective viruses. For this, medium was replaced with 500 µl inoculum. After 1 h incubation at 37°C, inoculum was removed, cells were washed with PBS and finally incubated with 500 µl culture medium. At certain time points, cell culture supernatant was harvested and centrifuged at 4,000 rpm for 5 min to pellet floating cells and the cleared supernatant was frozen at -80°C. To quantify cell-associated virus, infected cells were detached with accutase, centrifuged at 4,000 rpm for 5 min and cell pellets were resuspended in 500 µl culture medium. Virus was released from cells with three freeze-thaw cycles followed by centrifugation at 4,000 rpm for 5 min. Resulting supernatant was used for titrations.

Virus titrations were uniformly carried out on Vero76 cells, which were seeded in 24 well plates at 100,000 cells/well. On the next day, the culture medium was removed and cells were infected with virus supernatant in 10 fold dilutions for 1 h at 37°C. Thereafter inoculum was removed and replaced with overlay medium (2 vol. culture medium mixed with 1 vol. 3% avicel; FMC, Philadelphia, PA, USA) [47]. After incubation for 2-4 days, depending on the virus, the medium containing avicel was removed, cells were washed two times with PBS and then fixated by using cold methanol for 15 min at -20°C. For visualization of plaques, cells were stained with crystal violet solution (1g crystal violet, 100 ml ethanol in a final volume of 500 ml water) followed by one wash with water.

### Infection of neural aggregates

Infection of neural aggregates was performed in 24 well plates in a volume of 500 µl CDMIII medium containing virus. Based on a mean surface area of 11-12 mm2 we estimated that there would be approximately 2-3,000 cells on the surface of the neural aggregates. Therefore, we chose to infect with 2,000 pfu of either McHV1 or CeHV2, reflecting a MOI 0.5-1. Inoculum was diluted in CDMIII medium. For mock control, neural aggregates were incubated with fresh CDMIII medium without virus. After incubation of 1 h in a cell culture incubator (37°C, 80% humidity, 5% CO2), inoculum was removed, neural aggregates were washed in 500 µl DMEM followed by addition 500 µl CDMIII medium. At defined time points post infection (1, 24, 48 and 72 h), culture supernatant was removed and replaced with fresh CDMIII medium and stored for subsequent virus titration. After titration, the cumulative titer was calculated for each neural aggregate for each time point.

### Microscopy

For McHV1 infected samples brightfield images were taken at 10x magnification on an Olympus IX70 using Cell^F software. For all other viruses brightfield images were taken at 10x magnification using the ESID detector of a Zeiss LSM800 microscope and ZEN software. Images were adjusted in ImageJ [48] to cover the same area.

### Immunohistochemistry

Neural aggregates (70-100 days old) were fixed in 4% paraformaldehyde (PFA) solution for 20 minutes and washed 3 times with DPBS. Each fixed neural aggregate was then embedded in 2% agarose liquified at 50°C in 2 mL reaction tubes. Then agarose was chilled on ice for 5 to 10 minutes, to allow the agarose to solidify. Embedded neural aggregates were transferred to 4% PFA in 2 mL reaction tubes, for a second fixation, and incubated overnight on a shaker. After three washes in DPBS, the neural aggregates were embedded in paraffin and sectioned at 3 µm.

For immunohistochemistry, neural aggregates were deparaffinized and rehydrated using xylol and progressively decreasing concentrations of ethanol. Antigen retrieval was performed by microwaving the sections in 10 mM sodium citrate buffer (pH 7.6) for 10 minutes. Endogenous peroxidase activity was inhibited by peroxidase blocking reagent. Anti-HSV1+2 polyclonal rabbit antibody (1:200) (NB120-9533; Novus Biologicals, Wiesbaden, Germany) was used for the detection of viral proteins in McHV2-, CeHV2-infected, and mock-treated neural aggregates, anti-βIII-tubulin monoclonal mouse antibody (1:50) (T8660; Sigma Aldrich, St. Louis, MO, USA) was used as neuron marker. Anti-Rabbit IgG isotype was used in control stainings. Detection of the primary antibodies was carried out using Envision FLEX/HRP secondary antibody (GV80011-2; DAKO, Hamburg, Germany). 3,3′-diaminobenzidine (DAB) chromogen was used as substrate for the HRP, and Mayer’s hemalum solution was used as counterstain. Images of sections were taken using Aperio CS2 Slide Scanner and analyzed using Aperio ImageScope (Leica, Wetzlar, Germany) software.

For immunofluorescence staining, deparaffinization and antigen retrieval steps were performed as described above. The neural aggregate sections were blocked in 1% BSA in DPBS for 20 minutes at room temperature. After washing three times in DPBS, the sections were incubated (1 hour at room temperature or overnight at 4°C) with Anti-HSV1+2 (1:200) and anti-βIII-tubulin (1:50) antibodies. Subsequently and after washing with PBS, secondary antibody incubation (1 hour) was performed using AlexaFluor488™ goat anti-mouse IgG (Invitrogen, 1829920) (1:1,000) and AlexaFluor555™ donkey anti-rabbit IgG (Invitrogen, 2180682) (1:1,000). Incubation with DAPI (10 min, room temperature) (0.1 µg/mL) was used for nuclear stain. Stained sections were imaged using Zeiss Observer Z1 inverted fluorescence microscope and analyzed using ImageJ software [48].

## Results

For a detailed and systematic analysis of replication of human and NHP simplexviruses we used two well-characterized human viruses, HSV1 and HSV2, as well as the primate simplexviruses McHV1, PaHV2 and CeHV2. Additionally, we chose a set of four cell lines; one human, one from African green monkey and two from macaque. The different cell lines were selected to match the virus species; additionally, all of them have been previously used in infection experiments with different primate simplexviruses.

In a first experiment, we performed one-step growth curves to gain information on the replication kinetics. Vero76 cells, which were derived from an African green monkey, were used as positive control, since all viruses tested are routinely propagated in these cells [5, 9, 10, 25, 28, 29, 49]. In addition, we used the human A549 cell line, which has been used for virus isolation and functional studies of HSV1 and HSV2 [50-53] and was reported to be permissive to CeHV2 infection [54]. Finally, we employed two cell lines from rhesus macaques, LLC-MK2 [37] and TeloRF [39]. LLC-MK2 cells were previously reported to support replication of CeHV2 [55], while a CeHV2 reporter virus generated by us hardly grew in this cell line as well as in TeloRF cells [54]. To monitor virus replication, supernatants and cells from infected cultures were harvested over the course of 72 hours and virus titers determined by plaque assay.

Vero76 and A549 cells supported efficient production of all five simplexviruses, regardless whether supernatant or cell-associated virus was analyzed (Fig. 1 A, B). The highest titers were measured for McHV1 and HSV1, while CeHV2 titers were still increasing at 72 hours post infection (hpi). Infection of the two rhesus macaque cell lines, LLC-MK2 and TeloRF, resulted in two distinct patterns. Both cell lines supported efficient replication of McHV1 and PaHV2, while replication of HSV1 and HSV2 was inefficient on both cell lines (Fig. 1). CeHV2 replicated poorly on LLC-MK2 cells, while replication on TeloRF was detectable but did not reach the high levels of PaHV2 and McHV1. Finally, titers determined from virus-containing culture supernatants and cell-associated viruses were comparable. Thus, the simplexviruses studied could be grouped into viruses which replicated well on cells from rhesus macaques and those which did not.

**Figure 1.**
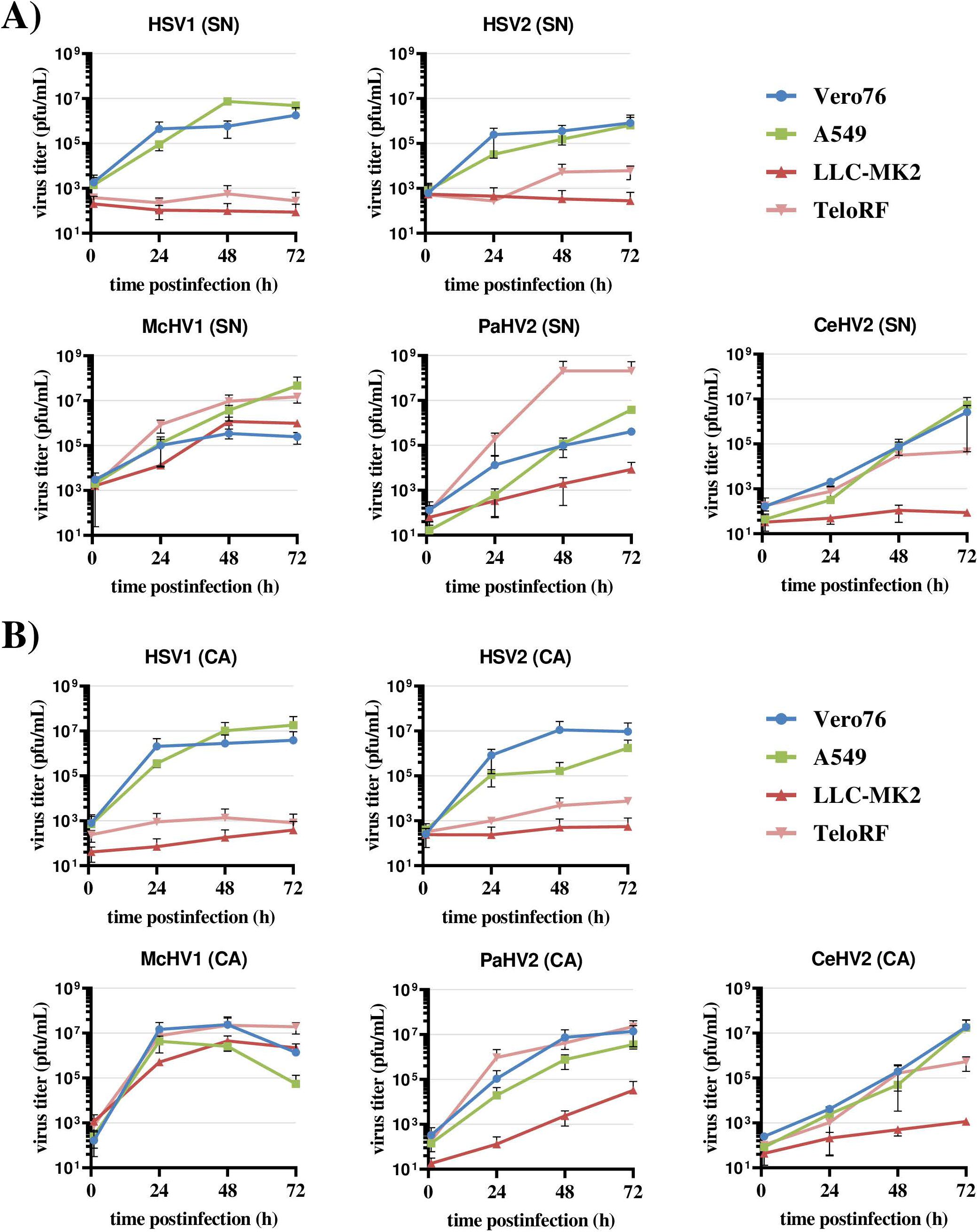
Comparison of primate simplexvirus replication in a panel of primate cells lines. Cell lines of human (A549), rhesus macacque (LLC-MK2, TeloRF) and African green monkey (Vero76) origin were infected with five primate simplexviruses at MOI1: Herpes simplex viruses type 1 (HSV1) and type 2 (HSV2), Macacine alphaherpesvirus 1 (McHV1), Papiine alphaherpesvirus 2 (PaHV2) and Cercopithecine alphaherpesvirus 2 (CeHV2). Virus titers from supernatant (SN, panel A) or infected cells (CA, cell associated; panel B) harvested at the indicated time points were determined by plaque assay on Vero76 cells. The results of two experiments carried out with triplicate samples are shown. Error bars indicate standard error of the mean, SEM.

All cell lines were monitored for cytopathic effects (CPE) throughout infection. Cell rounding and syncytia formation were detected as early as 24 hpi and increased up to 72 hpi (Fig. 2). The extent of syncytia formation differed between simplexvirus species. McHV1 and HSV2 produced large and prominent syncytia (Fig. 2 E-L). The capacity of PaHV2 to form syncytia was similar to that of McHV1 and HSV2 (Fig. 2 M-P). In contrast, CeHV2 and especially HSV1 induced syncytia formation with low efficiency (Fig. 2 A-D, Fig. 2 Q-T). In cell lines producing low titers of HSV1 or HSV2 (Fig. 3), no or minor signs of CPE were detected (e.g. Fig. 2 C, D, G, H). If virus was produced it appeared to originate from focal infection events, rather than widespread low-level replication.

**Figure 2.**
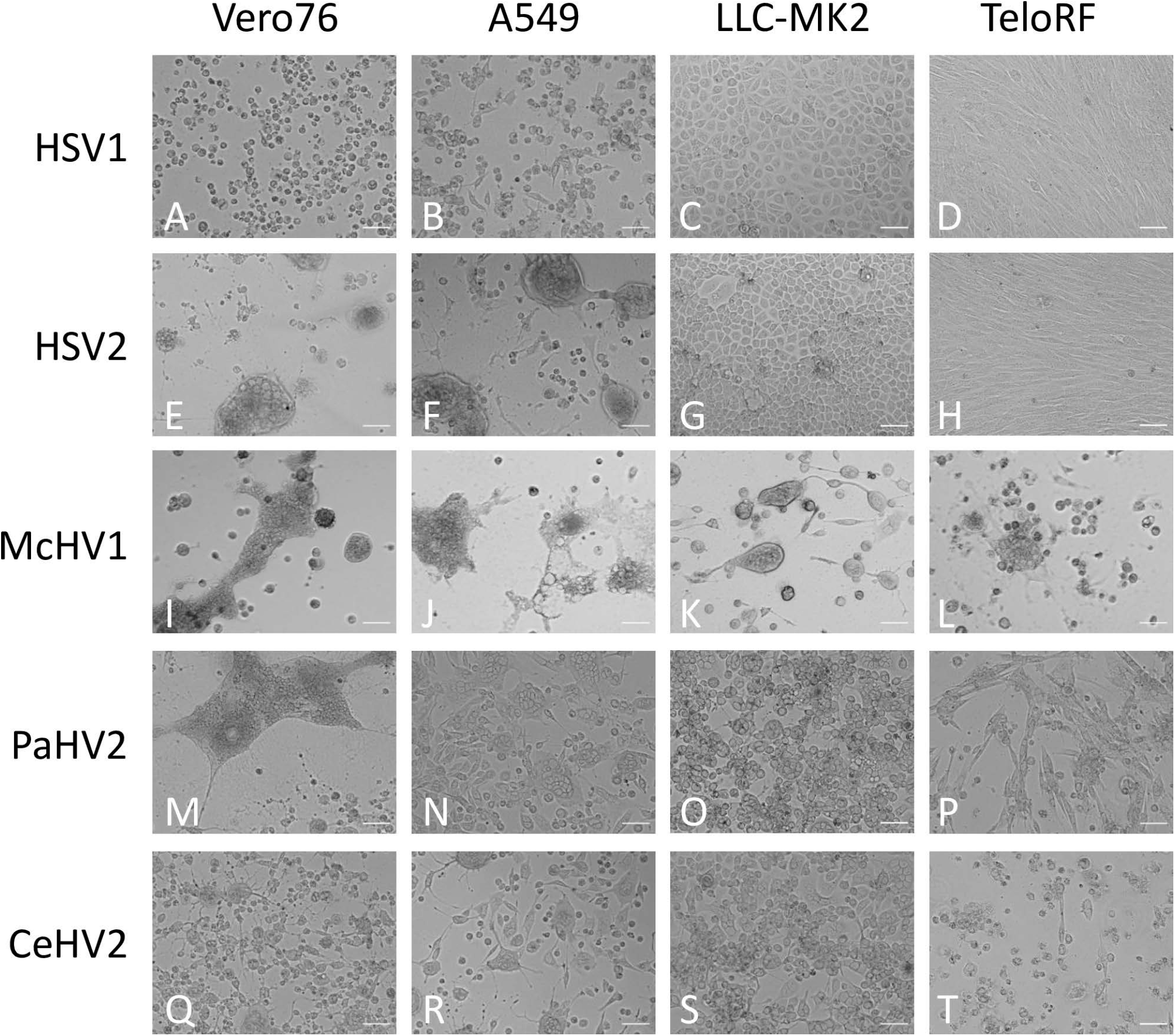
Cell morphology of infected cells. African green monkey Vero76, human A549 and rhesus LLC-MK2 and TeloRF cell lines were infected with five primate simplexviruses at MOI1. Brightfield images of infected cultures were taken at 72 hpi at 10x magnification. The scale bar indicates 100 µm.

**Figure 3.**
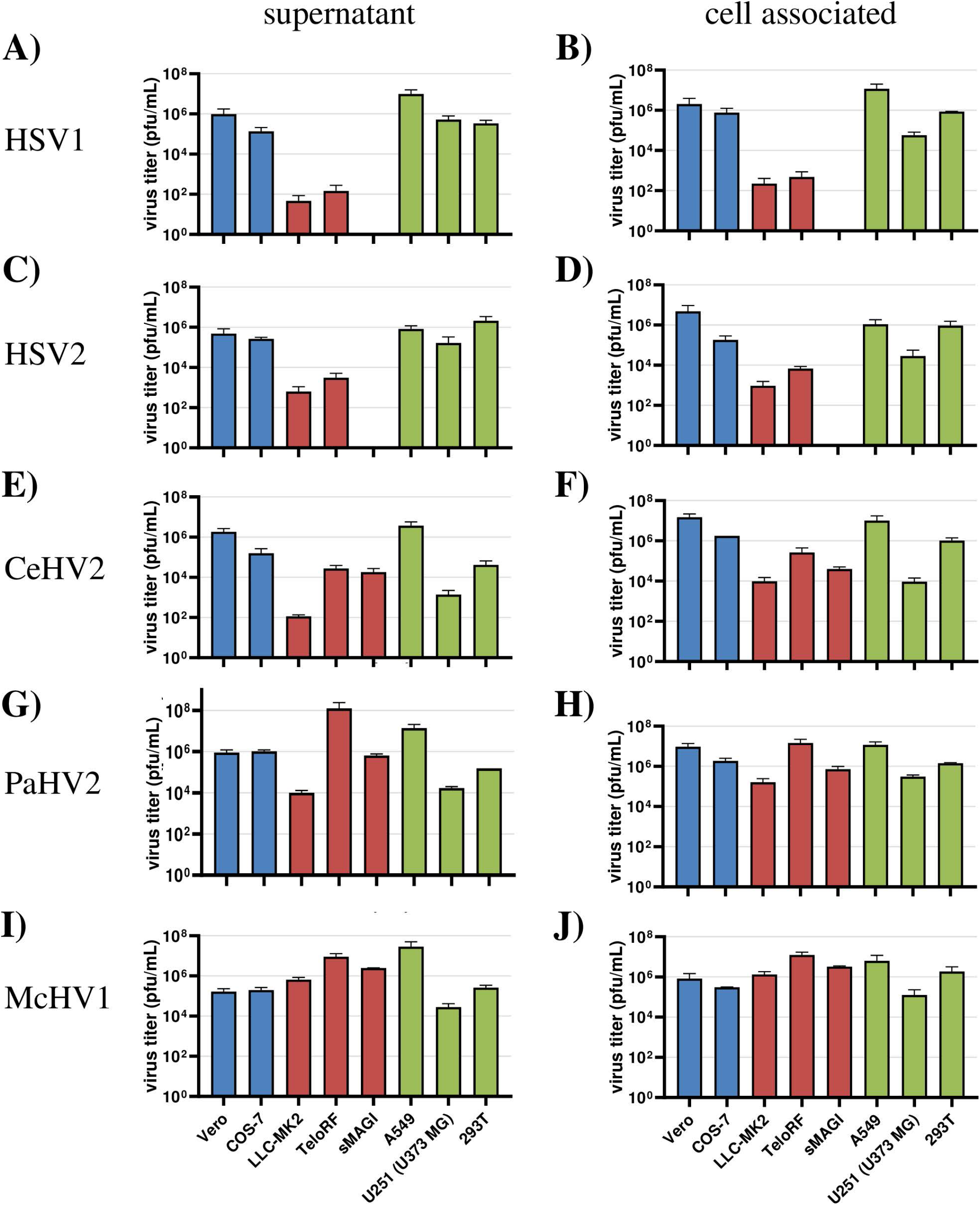
Comparison of virus production in a panel of primate cell lines. Herpesviruses HSV1 (A, B), HSV2 (C, D), CeHV1 (E, F), PaHV2 (G, H) and McHV1 (I, J) were used to infect human (A549, U-251 (U-373), 293T), rhesus macaque (LLC-MK2, TeloRF, sMAGI) and African green monkey (Vero76, Cos7) cell lines. Virus titers from supernatants (A, C, E, G, I) and infected cells (B, D, F, H, J) were determined by plaque assay on Vero76 cells. The results of two to four experiments carried out with triplicate samples are shown. Error bars indicate standard error of the mean, SEM.

To further extend our analysis, we expanded our panel of human cell lines, including the easy to manipulate 293T cells and the glioblastoma line U-251 (U-373 MG), which both are established in simplexvirus research [56, 57]. In addition, we used the African green monkey-derived Cos-7 cells and one additional epithelial cell line from rhesus macaques, sMAGI, as additional NHP cell lines. In this experiment, analysis was performed at a single time point postinfection, since our previous experiment revealed that the titers of most viruses reached their plateau at this time point independently of the cell line used. In Figure 3, we observed reduced replication of HSV1, HSV2 and CeHV2 in cell lines of rhesus macaque origin (LLC-MK2, TeloRF and sMAGI), while McHV1 and PaHV2 grew to high titers in all cell lines tested, with the only exception of PaHV2 showing reduced titers in the supernatant, when grown on LLC-MK2 cells. Furthermore, virus titers measured for supernatants (Fig. 3 A,C,E,G,I) were largely comparable to those of cell-associated viruses (Fig. 3 B,D,F,H,J), with the exception of HSV2, which repeatedly was not detected in the supernatant of infected TeloRF cells although low levels of cell-associated virus were detected (Fig. 3 C, D).

Finally, to mimic the pathophysiology of simplexvirus infection of the brain, we infected neuronal cells in a rhesus macaque 3D neural aggregate model. The model was based on rhesus macaque induced pluripotent stem cells (iPSCs). The 3D differentiation protocol was established according to published reports for human brain organoid generation [45, 46] (Fig. 4A). Successful neural induction was assessed in the neural aggregates after 70-100 days of differentiation by staining with general markers for neurons and glia cells (Fig. 4B). The aggregates contained neuron- and glia-like cells, assessed by immunostaining for cell-specific markers TUJ1 (for neurons) and GFAP (for glial cells), respectively (Fig. 4B).

**Figure 4.**
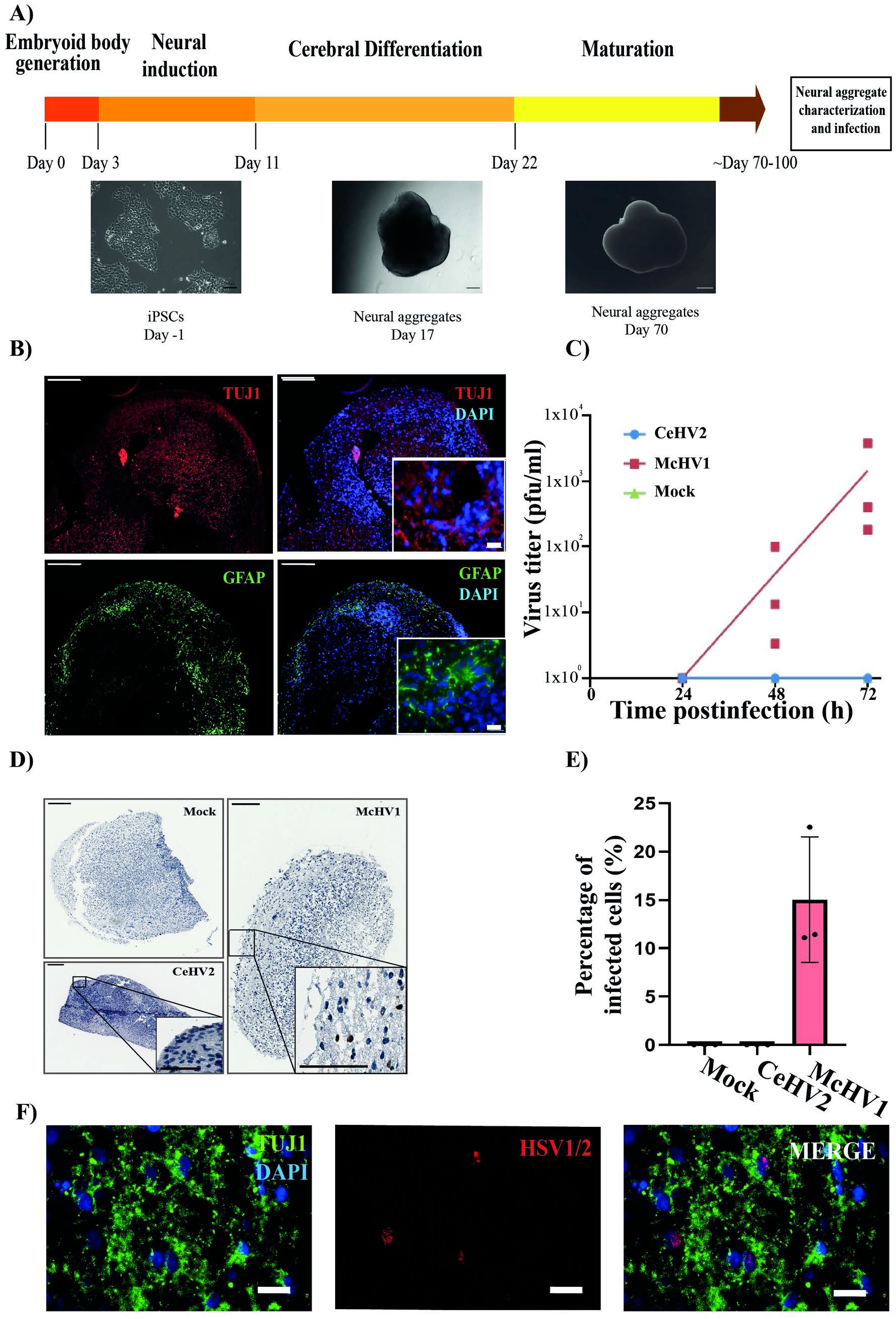
Comparison of McHV1 and CeHV2 infection of rhesus macaque neural cell aggregates. (A) Schematic representation of the neural aggregate generation protocol. Neural aggregates were derived from rhesus macaque iPSCs, and cultured for up to 100 days. Representative images of neural aggregates on day 17 and day 70 are shown. Scale bars: 100 µm, 200 µm, and 500 µm, respectively. (B) Neural aggregates were positive for a neuronal (TUJ1) and a glial marker (GFAP) at day 70. Scale bars: 200 µm; inset, 20 µm. (C) Rhesus macaque neural aggregates were infected in triplicates with 2000 pfu of McHV1 or CeHV2, respectively, or mock treated. Virus titers from supernatants of neurospheres were titrated on Vero76 cells. Dots show titers for individual organoids, while lines show the mean of three organoids. The results were confirmed in an independent experiment. (D) Immunohistochemical staining of the mock treated and infected neural aggregates for virus marker HSV1/2. HSV1/2+ cells were only found in the peripheral area of McHV1-infected neural aggregates (inset images). (E) Quantification of infected cells in immunohistochemical stainings. For each data point, three randomly chosen areas from three neural aggregate sections were analyzed. The total percentage of infected cells was calculated from the total nuclei in each area versus the number of simplexvirus-positive nuclei; error bars indicate standard deviation, SD. (F) Fluorescence imaging of the McHV1-infected neural aggregates for TUJ1 (neuronal marker) and HSV1/2 depicting viral proteins localized in nuclei of TUJ1+ cells. Scale bars, D: 400 µm; inset 200 µm. E: 20 µm.

From the previously tested panel of virus we selected McHV1 and CeHV2 to be validated in the 3D neural model. As shown in Figure 4C, productive infection could be demonstrated for McHV1 but not for CeHV2. Immunohistochemistry staining indicated a uniform distribution of McHV1 infection in the outer layers of the neurospheres (Fig. 4D) and confirmed absence of CeHV2 infection (Fig. 4D). Quantification of the infected cells was performed by image analysis of stained neurospheres. At the time of harvest (72hpi) 10-20% of the cells in neurospheres were infected with McHV1, while no infected cells could be identified in neurospheres infected with CeHV2 (Fig. 4E). Fluorescence imaging of the McHV1-infected neural aggregates showed the presence of McHV1 proteins in the nuclei of TUJ1+ neurons within the neural aggregates (Fig. 4F). Thus, McHV1 but not CeHV2 seems to have a high capacity to infect rhesus macaque neural cells.

## Discussion

Simplexviruses exhibit a broad species tropism, being able to infect many mammalian species from mice to humans [18]. Thus, when we established reporter viruses for CeHV2 and tested replication in cell lines from different species it came as a surprise that only very limited virus production was detected in several cell lines derived from rhesus macaques [54]. Although an older report, using wildtype CeHV2, came to conflicting conclusions [55], several publications also reported limited or no replication of HSV1 and HSV2 in rhesus macaque cells [31-33]. However, a comparative analysis was lacking so far. Our comparison of five human and primate simplexviruses in cell lines of human and NHP (rhesus macaque and African green monkey) origin shows that HSV1, HSV2 and CeHV2 have a limited capacity to infect rhesus macaque cell lines. In contrast, PaHV2 and McHV1 infected these cells efficiently, a finding that confirms and extends previous studies [32, 58, 59]. Importantly, studies with a rhesus macaque neural cell aggregate model demonstrated that the reduced capacity of CeHV2 to infect rhesus macaque cells extended to neuronal cells grown in a 3D culture system. In contrast, McHV1 replicated efficiently in this cell system, in agreement with observations for HSV1 in human brain organoids [60, 61]. In sum, we observed a differential capacity of primate simplexviruses to infect rhesus macaque cells.

We note a minor difference between our present and previous findings. Using wildtype CeHV2 we did not observe the strongly reduced replication in rhesus macaque cell lines that we had previously recorded for a CeHV2 reporter virus [54]. These differences might be related to the fusion of ICP4 with a reporter gene, which leads to reduced virus production (about 1 log lower), most likely due to impaired ICP4 expression. Regardless of the reasons for this discrepancy, it should be noted that replication of CeHV2 was still reduced by 2-4 log compared to McHV1 and PaHV2, underlining the differences in the capacity of NHP herpesviruses to replicate in rhesus macaque cells.

In the last years, it became clear that propagation of herpesviruses in cell culture can lead to rapid adaption due to selection of preexisting variants. Thus, it was shown that syncytia forming viruses are favored and rapidly selected for in cell culture, while this phenotype did not have an influence on virus titer [62]. In fact, earlier studies suggested that HSV1 and HSV2 may become adapted to rhesus monkey cell lines upon continued propagation [33]. For HSV1 it has been shown that propagation on baby hamster kidney cells leads to an expanded tropism towards previously nonpermissive Chinese hamster ovary cells. However, the underlying molecular reason for these adaptions is not yet known [63]. All viruses in our study have been extensively passaged on Vero cells [5, 9, 10, 49] and likely have adapted to these cells. However, despite this common adaption these viruses show clear differences in their ability to infect cells derived from rhesus macaques. We are therefore convinced that differences in the tropism to macaque cells cannot be explained by adaption to Vero cells.

Entry into target cells by simplexviruses is mainly mediated by glycoprotein D (gD), which binds several cell surface receptors of which nectin-1 is broadly expressed [18, 64]. Residues critical for nectin-1 interaction in gD (Y38, D215, R222, F223) are conserved among all herpesviruses used in this study [64]. Since McHV1 is not able to interact with Herpesvirus entry mediator (HVEM/TNFRSF14) [65, 66] and amino acid sequence among primate simplexviruses gD proteins is well conserved in regions required for HVEM interaction, it is likely that primate simplexviruses use nectin-1 as the main receptor for cell entry. In addition, glycoproteins of McHV1 and CeHV2 were well interchangeable in cell fusion assays [65]. This makes it unlikely that differences in receptor interaction are responsible for the differential entry into cell lines and neurospheres observed in this study.

The nature of the block to efficient infection of rhesus macaque cell lines with HSV1, HSV2 and CeHV2 remains to be elucidated. One can speculate that restriction factors of the innate immune system might be responsible. In fact, TRIM5α of rhesus macaque origin has been reported to reduce infection by HSV1 and HSV2 [67]. However, similar effects were also reported for African green monkey TRIM5α, making TRIM5α an unlikely candidate to explain the relative resistance of rhesus macaque cells as compared to human or African green monkey cells. Thus, it is likely that an as yet unknown factor is responsible for the selective restriction of HSV1, HSV2 and CeHV2 in rhesus macaque cells.

McHV1 can cause severe disease in humans and requires handling in BSL3 laboratories in Germany and BSL4 laboratories in the US. In contrast, CeHV2 and PaHV2 are believed to constitute a moderate threat to humans. While our study does not provide evidence that this concept should be changed, our finding that PaHV2 resembles McHV1 but not CeHV2 in its capacity to infect rhesus macaque cells might hint towards biological similarities between McHV1 and PaHV2. Indeed, for both McHV1 and PaHV1 neurovirulence in mice has been demonstrated, while CeHV2 was avirulent [21, 22, 68]. Additional comparative analyses are therefore needed to evaluate if this might be relevant for risk assessment.

## Acknowledgments

We would like to thank Wali Hafezi, Christiane Stahl-Hennig, David Brown, Matthew Jones, Thomas Stamminger, Sebastian Voigt and Andrea Maisner for their kind gift of material and Nicole Umland for excellent technical assistance.

